# Investigating the effects of molecular crowding on the kinetics of protein aggregation

**DOI:** 10.1101/2020.08.05.238584

**Authors:** John S. Schreck, John Bridstrup, Jian-Min Yuan

## Abstract

The thermodynamics and kinetics of protein folding and protein aggregation *in vivo* are of great importance in numerous scientific areas including fundamental biophysics research, nanotechnology, and medicine. However, these processes remain poorly understood in both *in vivo* and *in vitro* systems. Here we extend an established model for protein aggregation that is based on the kinetic equations for the moments of the polymer size distribution by introducing macromolecular crowding particles into the model using scaled-particle and transition-state theories. The model predicts that the presence of crowders can either speed up, cause no change to, or slow down the progress of the aggregation compared to crowder-free solutions, in striking agreement with experimental results from nine different amyloid-forming proteins that utilized dextran as the crowder. These different dynamic effects of macromolecular crowding can be understood in terms of the change of excluded volume associated with each reaction step.

## I. INTRODUCTION

Protein self-assembly is an important process for the formation of natural polymers like actin and microtubules,^1,2^ but also for the formation of amyloids,^3,4^ culprits for many neurodegenerative diseases including Alzheimer’s, Parkinson’s, type-2 diabetes, and others.^5–9^ Decades of protein self-assembly studies reveal that the molecular reactions involved are complicated and some new ones are still being discovered, especially in the field of amyloid formation.^10–17^ Experimental investigations have shown that both the folding of the constituent monomer proteins as well as the aggregate assembly are determined by a complex energy landscape, where numerous routes can convert monomer proteins into distinct aggregated structures that may or may not have biological functions.^18^

Among the many theoretical methods to investigate the protein aggregation problem, the kinetic approaches^10,11,14,19–26^ often provide direct fits to experimental data and interpretation of the aggregation processes. Processes considered in these studies and others often include primary nucleation, monomer addition, monomer subtraction, fibril fragmentation, merging of oligomers, heterogeneous (or surface-catalyzed) nucleation, etc. The reaction rates associated with the reaction steps considered can be independent or dependent on the oligomer/fibril size.^11,14,23^

Almost all of our current knowledge comes from the studies of systems *in vitro*, and little has been done in understanding processes *in vivo*. The inside of cellular environments is often crowded with proteins and other macromolecules or confined in compartments. In living cells, the volume fraction of the crowders can be as high as 40% of the total volume,^27–33^ that can include many different species of biological matter including biopolymers such as RNA, DNA, and proteins, organelles, metabolites, and osmolytes, resulting in a highly complex and packed environment. Confinement or volume effects on protein aggregation have received some attention recently, but the effects of molecular crowding on protein aggregation have so far been studied using only simple models,^34–36^ such as that introduced by Oosawa and Asakura^37^ or the Becker-Döring model.^38,39^

In the present article, we present a general kinetic study of the effects of molecular crowders on protein self-assembly using the kinetic equations of *P*(*t*) and *M*(*t*),^10,11,14^ which are the first two moments of the cluster size distributions. Previous treatments of the effects of molecular crowding on protein aggregation are reviewed in Section II. We introduce the kinetic processes considered in our treatment in Section III. The *P*(*t*) and *M*(*t*) kinetic equations without crowders are given in Sections III and IV and the changes of these equations due to the presence of molecular crowders presented in Section V. We then compare the results of numerical solutions of these kinetic equations to experimental data on nine different amyloid proteins in Section VI. Finally, some discussion and comments on our treatments are given in Section VII.

## II. PREVIOUS STUDIES OF THE EFFECTS OF MOLECULAR CROWDING ON PROTEIN AGGREGATION

Recent investigations probing the effects of crowding on protein aggregation have mainly focused on using molecular simulation to explicitly sample the interactions between proteins and crowders in 3D,^40–42^ and on using kinetic mass-action kinetics models for computing the changes to the cluster size distributions as a function of time.^23,35,36,43–45^ For example, Magno et al.^40^ used a simplified Lennard-Jones potential to quantify the interactions between the proteins and the crowders, and Langevin dynamics simulations to sample aggregation trajectories. They found that the crowders stabilized oligomers, and increased the solution viscosity. However, due to the expensive molecular simulations, they could only study very early states of aggregation. The mass-action kinetics models generally allow one to investigate the aggregation behavior at much longer time scales, but at a considerably less mechanistic detail into the assembly pathways.

Hall and Minton^35^ have studied the effects of macromolecular crowding on protein self-assembly using the Becker-Döring model^38,46^ as modified by Goldstein and Stryer,^43^ but for the *r*-th aggregate with *r ≤ n_c_*, the size of the critical nucleus, they assume the monomer-addition and -removal rates are size-dependent. However, for *r* greater than *n_c_*, they assume the rates are size-independent. Furthermore, they treat the (*n_c_* − 1) to *n_c_* reaction step differently to allow for a possible isomerization reaction leading to the formation of a critical nucleus.

The effects of molecular crowding are mainly taken care of through the change of the activity coefficients, which can be incorporated using transition-state theory for the monomer-addition reaction step. Two of the results that they have found are that the rate of fibril formation can be enhanced by orders of magnitude compared to crowderless solution and the degree of enhancement is strongly dependent on the size of the nucleus. Furthermore, they have also studied the corresponding equilibrium systems using similar but slightly different models.^34^

Bridstrup and Yuan have also studied the equilibrium systems of protein aggregation based on the Becker-Döring model.^36^ Similar to Hall and Minton, the effects of molecular crowding is treated based on the assumption that transition-state theory is valid. They have applied their method to fit the experimental data for actin with dextran or trimethylamine N-oxide^47^ (TMAO) as crowding agents as well as human apolipoprotein C-II with dextran.^48^ They have also studied the effects of molecular crowding on the kinetics of protein aggregation based on the Oosawa model. Their results fit the experimental data of Rosin, et al.^47^ of actin in the presence of dextran reasonably well. Furthermore, Minton and Hoppe investigated the effects of time-dependent crowding^44^ and surface adsorption^45^ on protein fibril formation.

## III. MICROSCOPIC REACTION STEPS INVOLVED IN AMYLOID FORMATION

In terms of the mass-action rate equations, we can write down the following microscopic reaction steps often involved in protein aggregation processes,^11,14,49^

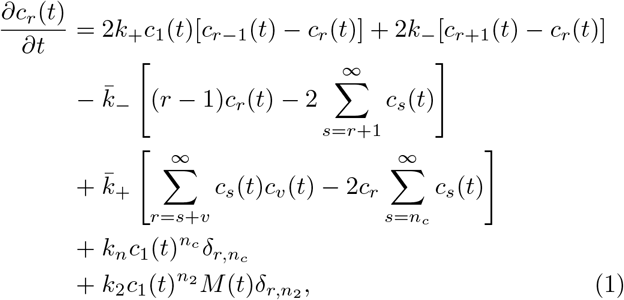

where *c_r_*(*t*) is the concentration of the aggregate of size *r* (refered to as an *r*-mer) at time *t*. Each term in Eq. (1) is illustrated schematically in Fig. 1. On the right hand side of Eq. (1), the first and second terms describe the reactions of monomer addition and subtraction with rate constants, *k*_+_ and *k*_−_, respectively, which are shown in Fig. 1(iv). The third and fourth terms describe the reactions of fragmentation of aggregates into smaller oligomers and coagulation of oligomers into larger aggregates, respectively, which are shown in Fig. 1(v) with rate constants, 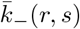 and 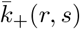. The fifth term describes the homogeneous nucleation with rate constant *k_n_*, which is shown in Fig. 1(i), and the sixth term describes a monomer-dependent 1-step secondary nucleation with rate constant, *k*_2_, which is shown in Fig. 1(ii). In the above equation, for simplicity, we assume that all rates are size-independent. The quantity *M*(*t*) is the polymer mass concentration defined by

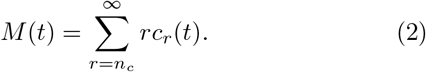

*M*(*t*) is the first moment associated with the size distribution, *c_r_*(*t*), whose zeroth moment defines the polymer number concentration, *P*(*t*), given by

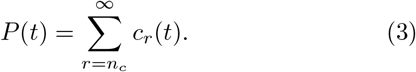

**FIG. 1.**
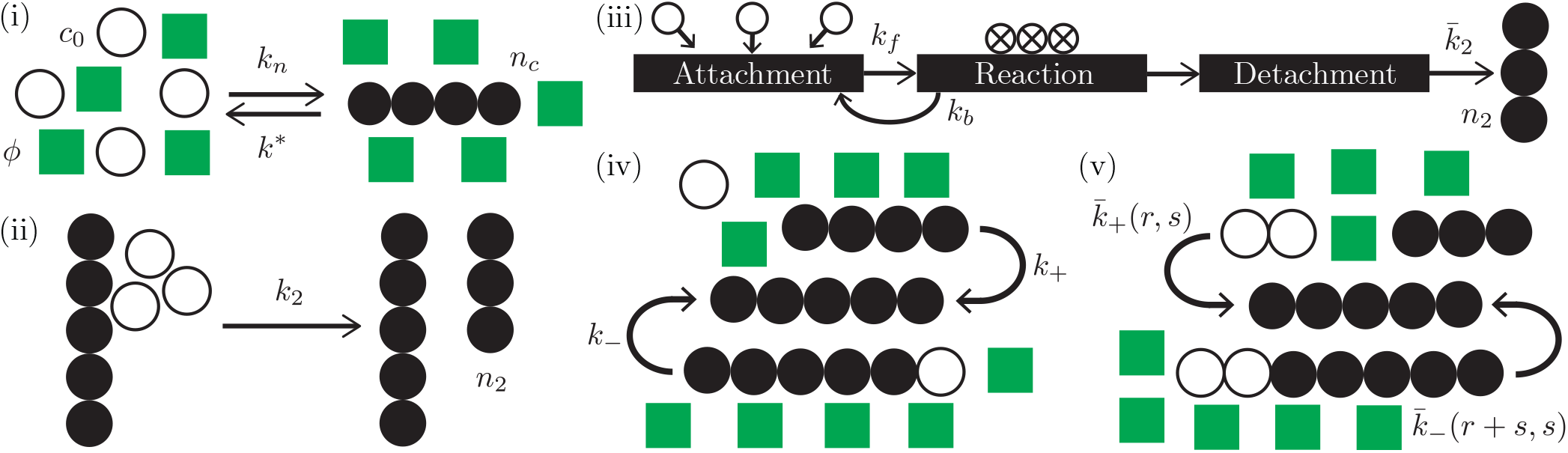
Schematic illustration of the different nucleation, growth, and shrinkage mechanisms described by Eq. (1). (i) Homogeneous nucleation of a cluster of size *n_c_*. The monomer-dependent secondary nucleation of a cluster of size *n*_2_ is illustrated in (ii) for 1-step nucleation and in (iii) for 2-step Michaelis-Menten-like nucleation. (iv) Monomer addition and subtraction mechanisms. (v) Fibril merging and breakage mechanisms, where the rate constants are shown as being size-dependent. In all panels, circles represent proteins and squares represent crowders. Open face circles show the proteins that are monomers either before or after the reaction, solid circles show proteins bound in a fibril, and circles in (iii) marked with an “X” show proteins bound to the surface of a fibril.

As such, the average length of fibrils, *L*(*t*), can be calculated as

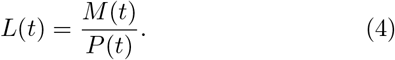

## IV. A CLOSED SET OF KINETIC EQUATIONS FOR THE MOMENTS OF THE SIZE DISTRIBUTION

We can study the molecular crowding effects on protein aggregation as done by Hall and Minton,^34,35^ and Bridstrup and Yuan^36^ by solving the mass-action equations like Eq. (1). However, a simpler and still accurate way to investigate the crowding effects is to use a closed set of the kinetic equations for the moments of the size distribution. Assuming rate constants are size-independent, a closed set of kinetic equations have been derived by Michaels and Knowles^11^ and many others.^14,50^ This is achieved by summing the mass-action equations, Eq. (1), over the aggregate size, *r*. We obtain

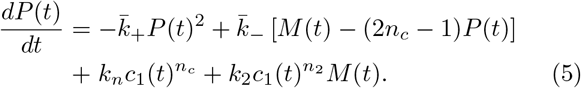

where we assume monomer detachment from a critical nucleus, 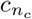, can be neglected. Similarly, we multiply the mass-action equations, Eq. (1), by size *r* and sum over *r*, to obtain

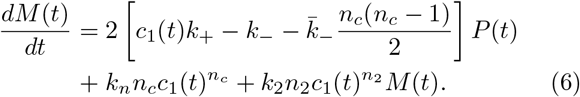

In the above equations, Eqs. (5) and (6), the monomer concentration, *c*_1_(*t*), is given by *c*_1_(*t*) = *c*_0_ − *M*(*t*), where *c*_0_ is the total monomer mass, which ordinarily is fixed.

## V. EFFECTS OF MOLECULAR CROWDING ON THE KINETIC EQUATIONS OF MOMENTS

In this section, we will work out the effects of molecular crowding on the kinetic equations of moments. In general, this is accomplished by modifying the rate constants in the kinetic equations describing protein aggregation. We begin by investigating first the general reaction steps in the mass-action rate equation where an *r*-mer and an *s*-mer combine into an (*r*+*s*)-mer and its reverse reaction, *M_r_*+*M_s_* ⇌ *M_r+s_*. The corresponding reaction rate constants are 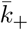 and 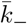, respectively, and in this reaction expression, *M_r_* denotes an aggregate made of *r* monomers. Consider, first, the forward association reaction, *M_r_* + *M_s_* → *M_r+s_*. The transition-state theory by Eyring^51^ assumes that a quasi-equilibrium is established between the reactants and transition states and the forward rate constant can be expressed as

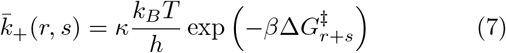

where *κ* is a constant, *k_B_* is the Boltzmann constant, *h* is the Planck constant, *β* is 1/(*k_B_T*), and

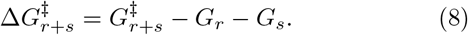

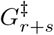 is the free energy of the transition state, and *G_r_* and *G_s_* are the free energies of an *r*-mer and an *s*-mer, respectively. At one mole we have

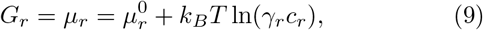

where *μ_r_* denotes the chemical potential and *γ_r_* the activity coefficient of an *r*-mer. Furthermore, the superscript 0 in 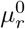 refers to the standard state. Using Eqs. (7), (8), and (9), the oligomer association rate constant is related to the ratio of the activity coefficients given by

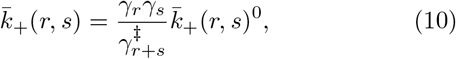

where 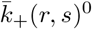 refers to the association rate constant in the case that all related activity coefficients are equal to unity. This includes, in particular, the case of a solution without the presence of crowder molecules. We shall call the latter the crowderless case below, when we investigate the effects of molecular crowders on reactions.

The effects of crowders on a reaction step can be investigated based on the changes of activity coefficients of reactants and products due to the presence of crowders in the solution. These coefficients can be calculated by the volume-exclusion method treating molecules as hard particles using the scaled-particle theory (SPT).^52^ We consider below how the oligomer association rate is modified by the presence of crowders. That means how crowders would affect the activity coefficient factor on the right-hand side of Eq. (10). The effects on activity coefficients depend on the shapes of the protein and crowder molecules. If we treat filamentous aggregates as sphero-cylinders, the expressions of the activity coefficients in the presence of crowders have been derived by Cotter using SPT.^34,44,53^ Here we introduce one more simplified assumption that the shape and geometry of the activated complex in the transition state closely resembles that of the product.^34,36,44^ We can then assume

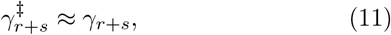

so that Eq. (10) becomes

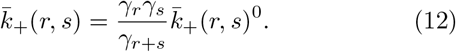

One key result from SPT for sphero-cylinders^34,36,44,53^ is that the activity coefficient for *r*-mer is related to that of a monomer through the following relation,

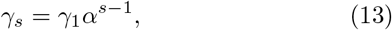

where *α* is parameter related to the excluded volume fraction, *ϕ*, of the crowders in solution, the ratio of the monomer radius *r*_1_ to the crowder radius *r_c_*, and the ratio of the sphero-cylinder radius, *r_sc_* to the crowder radius. It is defined in Bridstrup and Yuan.^36^ Using Eq. (13), we can then write the association rate constant with crowders present as^36^

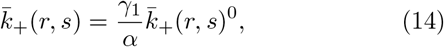

where the superscript 0 refers to the crowderless case. A similar calculation applies to the reversed reaction, fragmentation of an (*r* + *s*)-fibril. For this reversed reaction, the transition-state theory gives

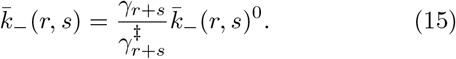

This time the assumption that the geometry of the activated complex resembles that of the complex, (*r* + *s*)-mer, as expressed in Eq. (11), gives the approximation,

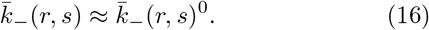

Of course, the important monomer-addition and subtraction reactions are just special cases of Eqs. (14,16). In these cases, these equations become, respectively,

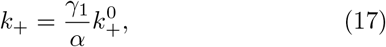

and

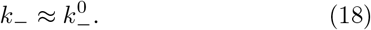

In the rest of the section we consider the effects of crowders on the homogeneous (or primary) and heterogeneous (or secondary) nucleation processes. First, for a primary nucleation process as shown in Fig. 1(i), we use again the transition-state theory to obtain

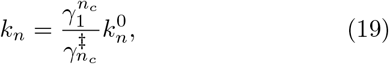

where 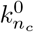 is the nucleation rate in a crowderless solution. Again the assumption that the geometry of the activated complex resembles that of the critical nucleus leads to

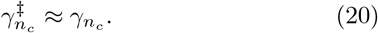

To proceed, we introduce a simplifying approximation that the nucleus shape resembles that of a sphero-cylinder, which may not be rigorously true in general. Under the sphero-cylinder assumption, we have^36^

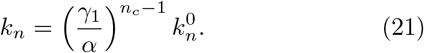

We next consider heterogeneous, or surface-catalyzed, nucleation processes. These processes can be classified into different types, for example, one-step or multi-step secondary nucleation processes, as illustrated in Fig. 1(ii) and (iii), respectively (see also Fig. 3 of Meisl, et al.^50^). The effects of molecular crowding will be different for different types of secondary nucleation processes considered. As presented in Eqs. (5) and (6), we have assumed that secondary nucleation is a one-step process (Fig. 1(ii)), which is accounted for by the last terms on the right-hand-side of these equations. The effects of crowders on the rate constants of these terms can obtained as a limiting case of a two-step secondary nucleation process. Thus, we first focus on a two-step, surface-catalyzed nucleation process, as is illustrated in Fig. 1(iii). A surface-catalyzed process can be considered a special case of an enzyme-catalyzed process and therefore can be treated with a Michaelis-Menten-like (MM) model.^54^ The crowderless version has been previously studied by Meisl, et al.^49^ In this description, at an active site on the surface of a fibril, *n*_2_ monomers are catalyzed to form an *n*_2_-mer, then this *n*_2_-mer goes into the solution as is illustrated schematically in Fig. 1(iii) (notice that this last step is different from the process considered by Ferrone, et al.,^55^ where a branch grows on the side of a polymer). According to the MM model, the catalyzed reaction can be described by the following two steps:

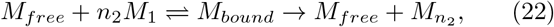

where *M_free_* describes a free site on the surface of a fibril on which *n*_2_ monomers can bind with a rate constant, *k_f_*, and *M_bound_* describes the corresponding bound complex. The reverse reaction of the first step has a rate constant, *k_b_*. The intermediate *M_bound_* can also proceed to produce an *n*_2_-nucleus in the solution, 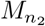, plus a free site with a rate constant, 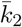. First, we examine how crowders would affect the formation rate of the bound complex, the first forward step of the MM model. Again, applying transition-state theory to this formation step leads to the following expression for *k_f_*

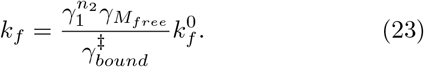

As before, based on shape similarity, we make the assumption that 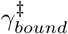 can be approximated by *γ_bound_*. Let us define

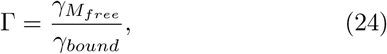

which can be evaluated in the following two approximate ways, introduced by Ferrone, et al.^55,56^ In the first approximation, we treat *M_free_* and the complex, *M_bound_*, as crystals and assume their activity coefficients to be unity^55^, thus Γ in

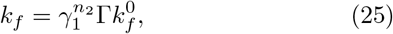

is just unity. The value of Γ can be greatly improved in the second approximation using a formula developed by Boublik.^57,58^ Following Ferrone^56^ and Minton,^58^ we obtain

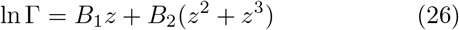

where

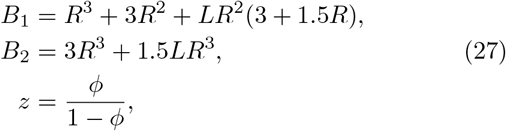

*R ≡ r_sc_/r_c_* and *L ≡ l*/2*r_sc_*. In these expressions, *ϕ* and *r_c_* are, respectively, the volume fraction and radius of crowder molecules, *l* is the cylindrical length of a nucleus (the portion between the hemispherical caps) and *r_sc_*, its radius. Conservation of volume between a nucleus and *n*_2_ monomers gives us

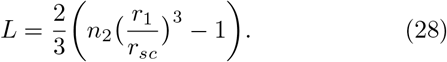

The MM-like model tells us that the heterogeneous nucleation contributes to the rate of generating a secondary nucleus, 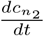, by a term given by

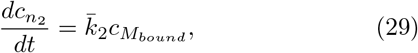

where 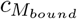 denotes the concentration of the surface-bound complex. Imposing the steady-state assumption, as is usually done in an MM model, we arrive at a term replacing the last term on the right-hand side (RHS) of 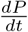 in Eq. (5) as follows:

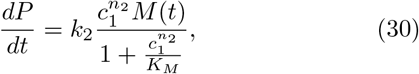

where *k*_2_ and *K_M_* are defined by

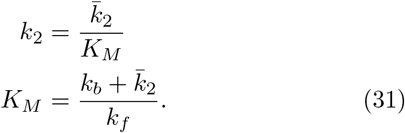

Similarly, the last term on the RHS of Eq. (6) should be changed accordingly, that is, replaced by the term on the RHS of Eq. (30), multiplied by *n*_2_. The reasoning that we have presented earlier about the negligible effects of crowders to the dissociation rate of the (*r* + *m*)-mer also applies here to *k_b_* and 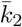. Appreciable crowding effects exist only in *k_f_*, but then also in *k*_2_ and *K_M_* as a consequence. However, in the limiting case when 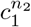 is much greater than *K_M_*, the crowding effects cancel and the heterogeneous nucleation rate is then not affected by the presence of crowders.

More interesting is the other limit, when 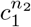 is much smaller than *K_M_*. Here we recover the form of the last term on the RHS of 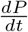 in Eq. (5), representing the one-step secondary nucleation process in the presence of crowders. This is because *k*_2_ is defined by Eq. (31), which contains the effect of crowding in *K_M_* through *k_f_*. The true crowderless limit is recovered when both activity coefficients, *γ*_1_ and Γ, approach unity in the absence of crowders. We note that in the presence or absence of crowders, the secondary nucleation term in 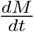, the last term on the RHS of Eq. (6), changes correspondingly, through *k*_2_, for the one-step process (Fig. 1(ii)).

### A. Changes to the solvent viscosity due to the presence of crowders

So far we have focused on the changes to the rate constants caused by changes to the entropy of the system due to the presence of crowders. Before solving the rate equations, we model the influence that the crowders have on the viscosity of the solvent, which SPT was not constructed to describe. Ioan, et al.^59^ points out that the friction of the solvent f(*ϕ*) relates to the hydrodynamic radius of the crowder, *R_h_*, through

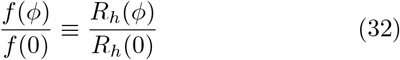

where the quantity is *R_h_*(*ϕ*) a correlation length that describes a long-range effect of frictional interaction among the dextran crowders.^59^ We treat *R_h_*(*ϕ*) in Eq. (32) as an “effective” radius for the crowder, which will grow with *ϕ*. Only for highly-dilute systems can this relationship be linear. In general *R*(*ϕ*) is a non-linear function and may change by up to an order of magnitude compared to the crowder-free radius for very high crowder concentrations.^59^ We therefore choose to model the effective crowder radius, 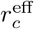, according to

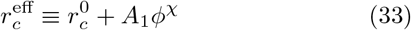

where 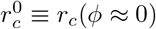 is the effective radii at low volume fraction, while the second term is a non-linear correction added to 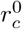. The parameters *A*_1_ and *χ* quantify the concentration dependence, and can be used to parameterize the model to experimental data, with *χ* intended to capture the leading order relationship between solvent friction and the crowders.

## VI. RESULTS

### A. Fitting the model to experimental data

The quantity *M*(*t*) is used as the fit function to be parameterized using experimental ThT data (see the Supporting Information for details on obtaining the half-times from experimental data, as well as other details). This procedure requires ThT measurements for several initial mass concentrations, from which the dependence of the half-time *t*_1/2_, on the initial monomer concentration, *c*_0_ can be determined. These two quantities are related via ln(*t*_1/2_) = *γ* ln(*c*_0_) + constant, where *γ* is the scaling exponent that is related to the reaction order of the dominant mechanism(s) (which allows one to determine *n_c_*). Any deviation from a straight line indicates that the dominant mechanism(s) have changed. All of the crowded protein aggregation experiments that we consider here had *c*_0_ fixed while varying *ϕ*. The reaction order (hence *n_c_* and *n*_2_) for the different proteins studied in experiments had to be determined from previous studies not involving crowders. We use the half-times obtained from the crowding experiments to probe the dependence of the dominant mechanisms on the presence of crowders.

To fit the ThT data for kinetics experiments involving crowders, the quantity *M*(*t*) is first fit to the crowder-free data to obtain the rate constants. In a previous study,^36^ we fit an effective crowder radius for each (*c*_0_, *ϕ*) set of ThT data while keeping the rate constants fixed, hence the number of additional parameters needed grew with the number of measurements made at different *ϕ*. Additionally, the protein monomer radius *r*_1_ and the radius of the spherocylinder assembly *r*_sc_ were assumed to be constant and were not used as fit parameters. Modeling the effective radius dependence on *ϕ* by means of Eq. (33) requires two additional parameters (*A*_1_ and *χ*) no matter the number of half-time measurements made at different *ϕ*. We first fit the model to experimental data with *r*_1_ = *r_sc_* = *r*, following Ellis and Minton^61^ by modelling the filaments as one-dimensional assemblies, where *r* is an additional fit parameter. If the model cannot be fit to the data under this assumption, we use both *r*_1_ and *r_sc_* as fit parameters. Hence, in total, up to four extra parameters are required to capture the influence of the crowders on the protein aggregation kinetics for different proteins.

We also showed previously that atomic force microscope (AFM) and ThT measurements can both be used to obtain more robust estimates of the rate constants.^23^ The ThT data sets for the crowder experiments studied here were not accompanied by AFM measurements. Nevertheless, we added a small bias to the objective function (the mean-average error between predicted and measured fibril mass concentrations) to encourage physically realistic rate constant parametrizations for fibril lengths (maximal *L*(*t*) values) in the range of hundreds to tens-of-thousands of proteins.^8,62,63^

### B. Comparison with experiments

We first consider actin self-assembly with dextran crowders present in solution to illustrate the fit procedure. The critical nucleus size was first determined to be *n_c_* = 3 by using ThT data from Rosin, et al.^47^ to obtain *γ*. Fig. 2(a) shows the experimentally measured ThT response (symbols) and the mass concentration *M*(*t*) (lines) computed using Eq. (1) for different crowder volume fractions *ϕ*. The results in Fig. 2(a) show that as *ϕ* increases, M(t) rises earlier and faster with time. The asymptotic value of M(t) also increases with *ϕ*. Fig. 2(b) shows the predicted length profiles at different *ϕ* with corresponding *r_c_*. Fig. 2(c) illustrates the dependence of *t*_1/2_ on *ϕ*, which looks like a decaying exponential. The solid line was computed by taking *r_c_* equal to the average of the fitted values for the *ϕ* > 0 data sets.

**FIG. 2.**
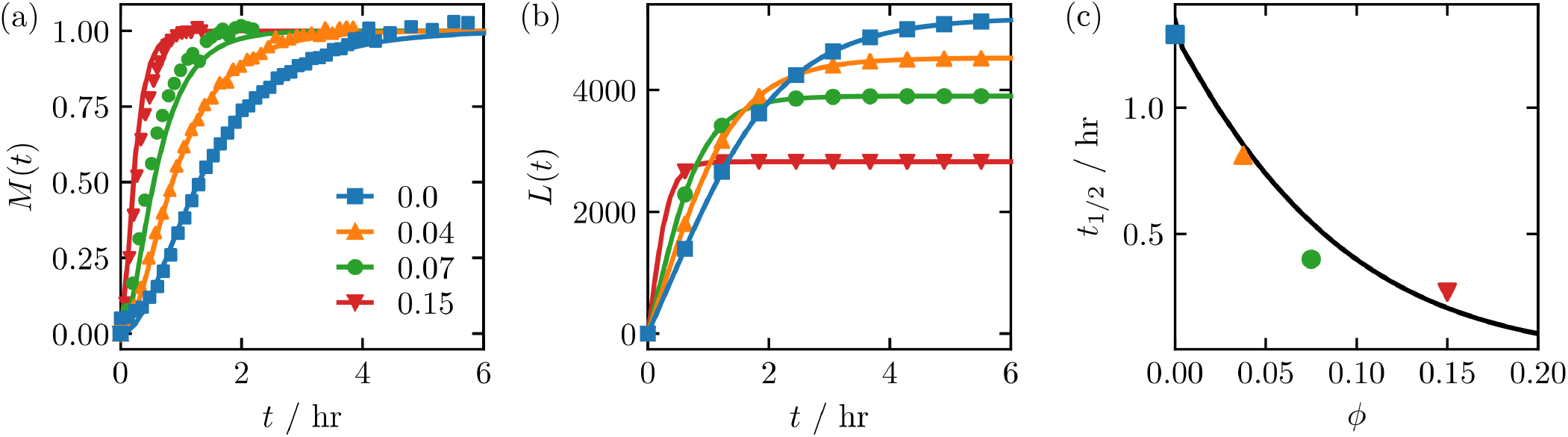
Actin. In (a) the mass concentration of fibrils *M*(*t*) is shown and was fit to ThT data from Rosin, et al.^47^ using a least-squares routine. In (b) the predicted *L*(*t*) is shown for different crowder concentrations *ϕ*. In (c) the half-time (*t*_1/2_) obtained from the curves in (a) is shown versus *ϕ*. Non-zero rate constants obtained from the fit are listed in the Supporting Information. The different symbol shapes indicate different values for *ϕ*. In (a) they also refer to experimental data.

Fig. 3 shows the scaling dependence on *ϕ* for various protein aggregation experiments.^47,48,60,64–68^ Additionally, plots of *M*(*t*) and *L*(*t*) for all proteins shown in Fig. 3 can be found in the Supporting Information. The proteins appear to fall into two groups. The first group (denoted group 1) responds to the presence of crowders by accelerating the aggregation (negative slope), and includes actin,^47^ human apolipoprotein C-II (ApoC2),^48^ human bovine prion protein (PrP (Hu)),^64^ beta-2-microglobulin (*β*2m),^65^ beta-lactoglobulin (*β*-LAC),^66^, and amyloid-beta-peptide 1-40 (*Aβ*(1 − 40)).^60^ The second group (denoted group 2) shows a slow down of the aggregation (nearly flat, or positive slope), and includes the human islet amyloid polypeptide (IAPP),^67^ rabbit bovine prion protein (PrP (Ra)),^64^ and bovine carbonic anhydrase protein (BCA).^68^ Numerous other protein aggregation experiments involving crowders have observed similar behavior.^69^ The large range of *t*_1/2_ is determined by the relative magnitudes of the different rate constants that are specific to each protein.

**FIG. 3.**
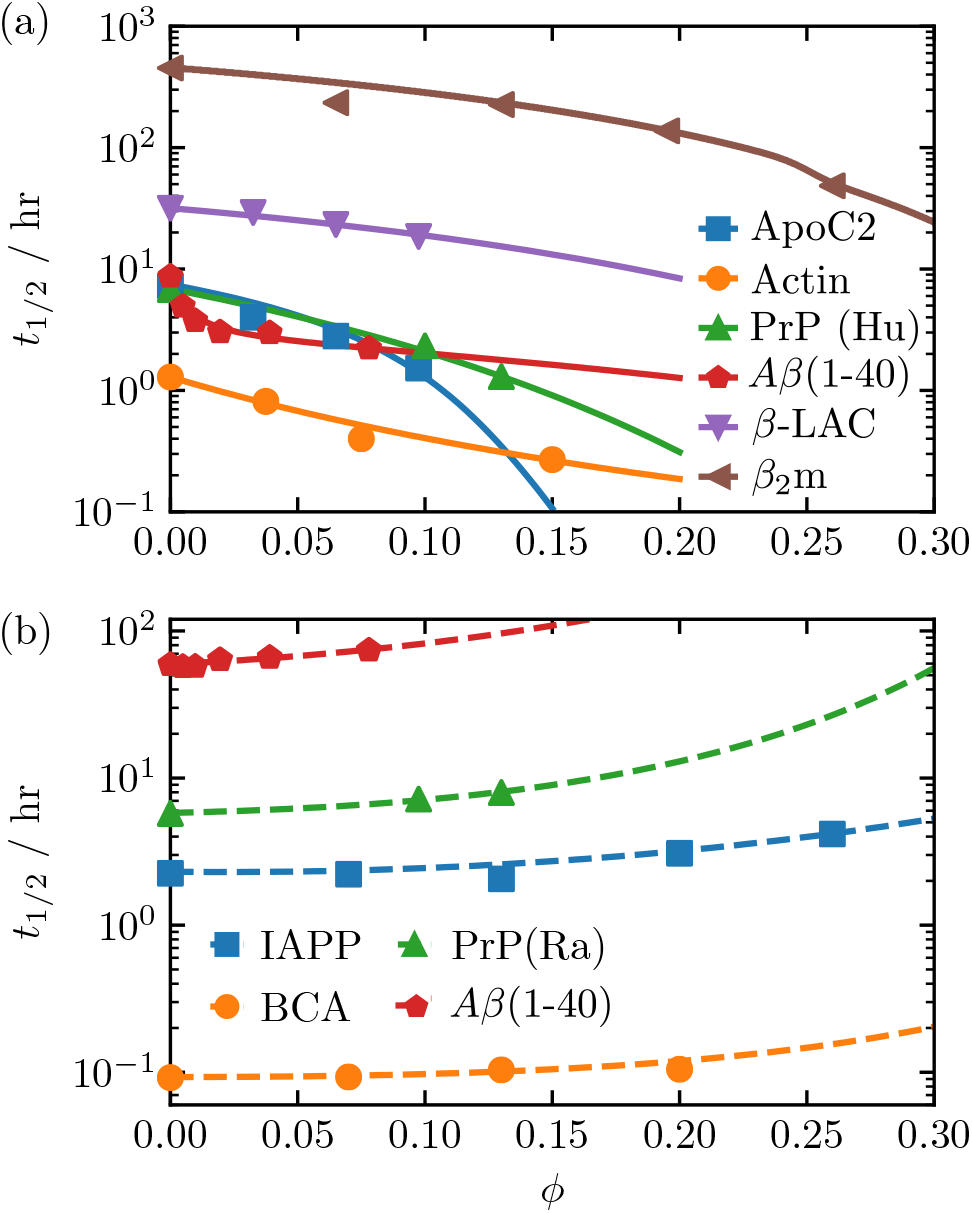
The half-time dependency on *ϕ* is shown for several different proteins. Experimental and computed values are shown with markers and lines, respectively. Panel (a) and (b) show proteins in which the half-times increase (solid lines) and those where it remains flat or decrease as a function of excluded volume, respectively. The A*β*(1-40) data in (a) was obtained in an agitated (shaken) environment, while that in (b) was obtained without agitation.^60^ All fit parameters are listed in the Supporting Information.

The aggregation of proteins in group 1 conform to the ordinary concept that is expected based on SPT– the cause for the aggregation speed-up is purely entropic in nature. For an association or merging reaction, the excluded volume decreases as proteins or oligomers merge into a larger aggregate, if *r_sc_ ≥ r*_1_. The reduction of excluded volume implies that the number of configurations in the fluid for the crowders increases, that is, the entropy increases, for the forward reaction. Thus, the second law of thermodynamics favors the association reaction, which increases its rate in the presence of crowders. Fig. 3 shows that SPT predicts that the dependency of *t*_1/2_ on *ϕ* can be either concave up (A*β*, actin) or concave down (*β*-LAC, *β*2m, PrP, ApoC2), in accordance with the experimental observations.

Figures 3(b) and 4 further show that SPT predicts that the aggregation for some proteins is slowed by the presence of crowders. For the four proteins in Fig. 3(b), we found that *r_sc_ < r*_1_ through fitting. Fig. 4, which shows the quantity *γ*_1_*/α* versus *ϕ*, which controls the change to the association and nucleation rate constants in Eqs. (17) and (21), clearly shows a ratio *r*_1_ ≤ *r_sc_* ≈ 0.8 for which crowders essentially do not affect the growth rates (red line) for increasing *ϕ*. Values greater than 0.8 show that the aggregation speeds up with increasing *ϕ*, while the opposite is observed for ratios below 0.8 where aggregation is increasingly slowed down with increasing *ϕ*.

**FIG. 4.**
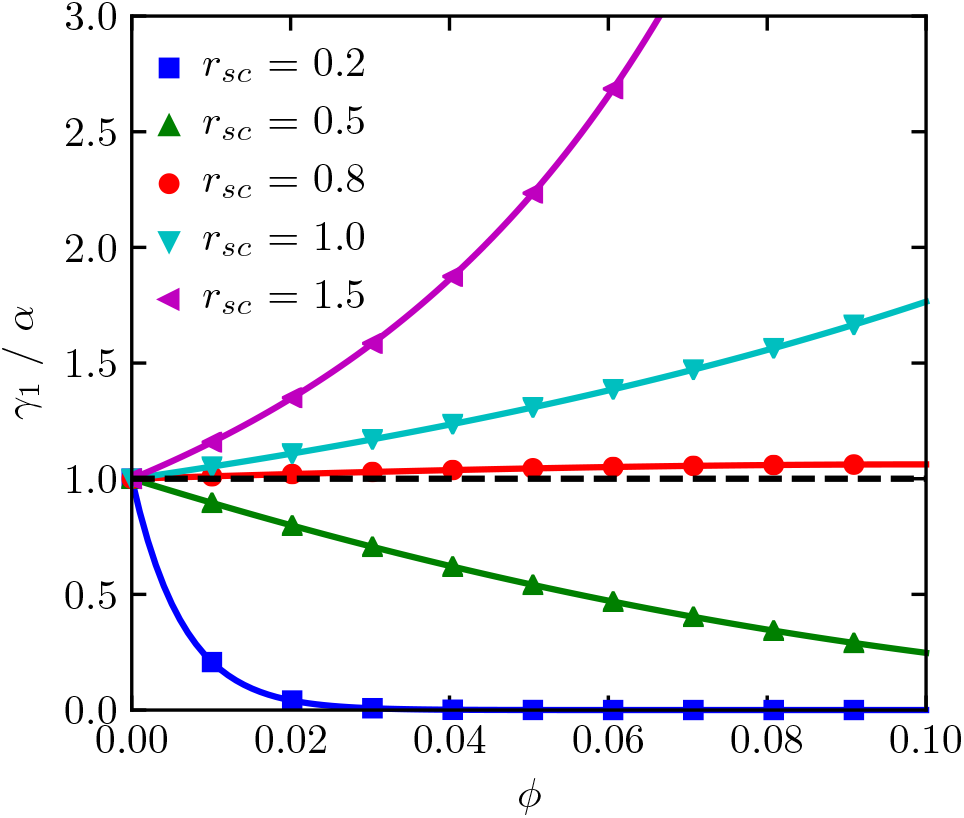
The quantity *γ*_1_/*α* is plotted against the excluded volume fraction *ϕ*. The different curves show the quantity computed for different values of *r_sc_* with *r*_1_ = *r_c_* = 1 nm. The dashed horizontal line shows *γ*_1_/*α* = 1.

The reason for the slow down is still due to changes in the entropy or excluded volume in the system. In some limit of the systems which satisfy the condition *r_sc_* < *r*_1_, the excluded volume can increase for the association reaction. For example, the aggregate formed can be long and thin (the assumption is that the volume is conserved when the sphero-cylinder of the aggregate is formed.^34^) Based on the same thermodynamic reasoning, the rate of association would slow down for this case. In Section VII we discuss these results in further detail.

### C. Crowders influence *L*(*t*) profiles

For the actin data set, the rate constants obtained from the fit in Fig. 2(a) tell us that homogeneous nucleation initiates the aggregation by converting monomers into critical nuclei that then elongate via monomer addition. Fig. 2(b) shows the computed *L*(*t*) increasing monotonically and then converging to a fixed value within the time-scale of the experiments (~6 hours). For *ϕ* > 0, Fig. 2(a) shows the proteins binding to the ThT dye at earlier times, clearly indicating a speed-up in the aggregation. Fig. 2(b) also shows that *L*(*t*) initially increases more rapidly as *ϕ* increases, but then plateaus at earlier times and at shorter lengths. Since no shrinkage mechanisms were found to be important, this result arises from the fact that larger *ϕ* corresponds to greater speed-up of the primary nucleation process, which generates more active clusters. Since the total number of monomers is constant, this results in comparatively shorter fibril lengths at long experimental times compared with lower *ϕ*.

In Fig. 5(a), *L*(*t*) is shown again for actin using the rate constants obtained from Fig. 2(a), except for *k_−_*, which is varied. As *k_−_* increases from zero, the fibril lengths remain unchanged from that shown in Fig. 2(b). Once monomer subtraction switches on (Fig. 5(a)(ii)), fibril lengths become reduced. Note that the effective increase in protein concentration due to the presence of the crowders mitigates the influence of monomer shrinkage compared to the crowder-free case. This effect is seen more clearly in Fig. 5(a)(iii) when monomer shrinkage is strong.

**FIG. 5.**
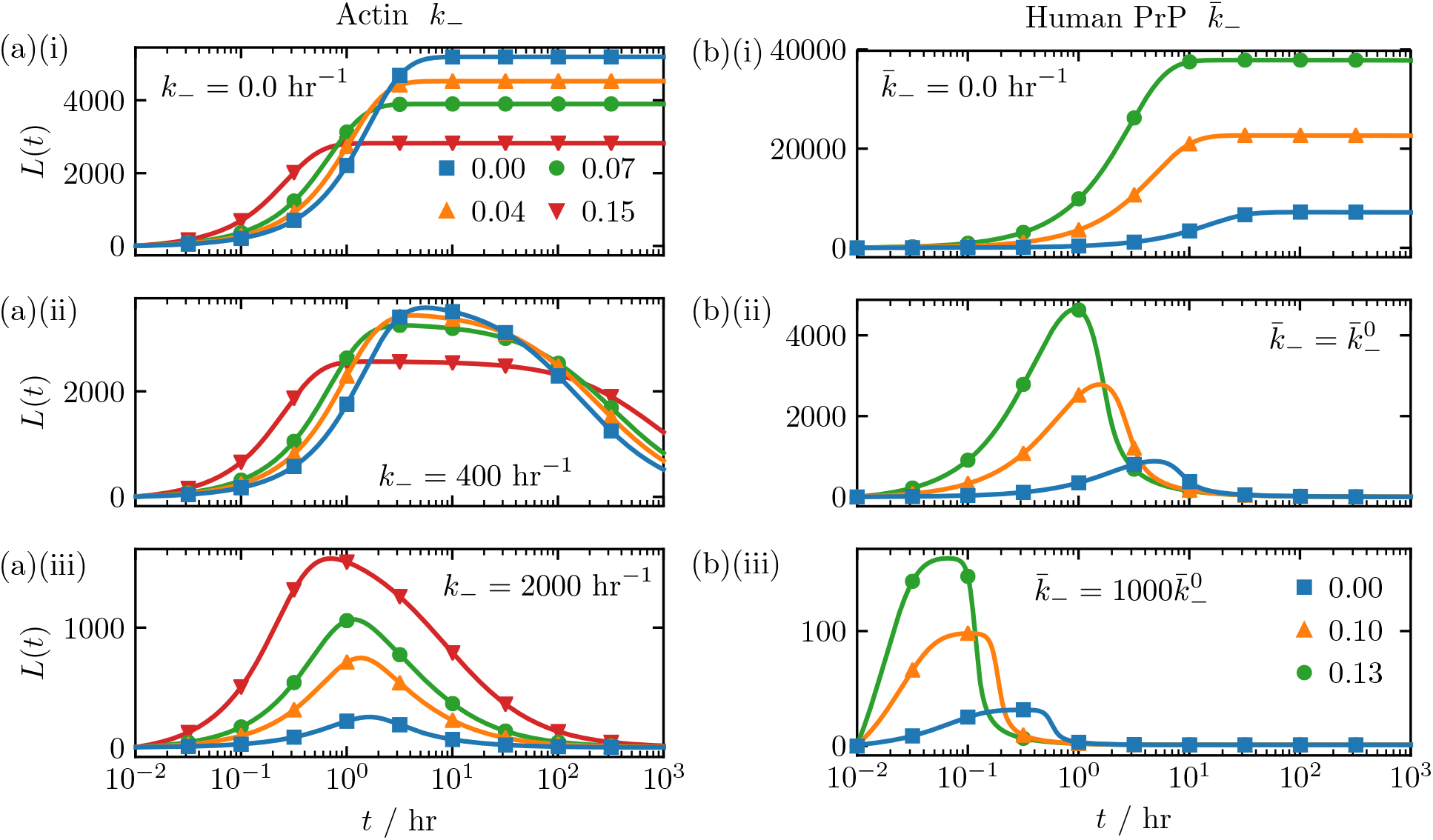
Length profiles for actin and human PrP are shown in (a) and (b), respectively. Unless otherwise noted in panels (i-iii), the fit parameters for these proteins are listed in the Supporting Information.

For the other protein systems shown in Fig. 3, the fibril lengths have a non-trivial dependence on *ϕ*. For example, *L*(*t*) is shown in Fig. 5(b) for human PrP. The mechanisms affecting the aggregation that were determined from fitting *M*(*t*)/*c*_0_ to the ThT data at *ϕ* = 0 are monomer addition and subtraction, fibril fragmentation, and a nucleus size *n_c_* = 1. Fig. 5(b) shows *L*(*t*) for several values of 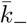. The length profiles clearly contrast those for actin. For example, Fig. 5(b)(i) shows that the lengths grow longer with increasing *ϕ* and reach a plateau within 10-100 hours. As fragmentation becomes relevant and 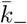 grows larger (Fig. 5(b)(ii) and (b)(iii)) the fibrils are both increasingly shorter and disintegrate at earlier times as *ϕ* increases.

Crowders may also influence the length profiles by affecting fibril elongation mechanisms, namely monomer addition, and fibril merging. For example, Fig. 6 illustrates the changes to *L*(*t*) for the ApoC2 protein, when the relative importance of the elongation mechanisms are considered. Monomer subtraction, fibril disintegration, and *n_c_* = 5 also influence *L*(*t*). When fibril merging dominates elongation, (Fig. 6(a)) increasing *ϕ* causes the fibrils to grow increasingly longer as fibrils merge together at a higher rate. Fig. 6(b) shows the scenario when monomer addition is competitive with merging. At early times, critical nuclei are initially scarce but elongate quickly, initially overshooting a stable (steady-state) value. As aggregate numbers increase and monomers become scarce, fragmentation takes over and breaks up the fibrils, whose average length approaches a steady-state value at longer times. Increasing *ϕ* mitigates the shrinking phase somewhat because the higher fibril merging rate counterbalances fragmentation.

**FIG. 6.**
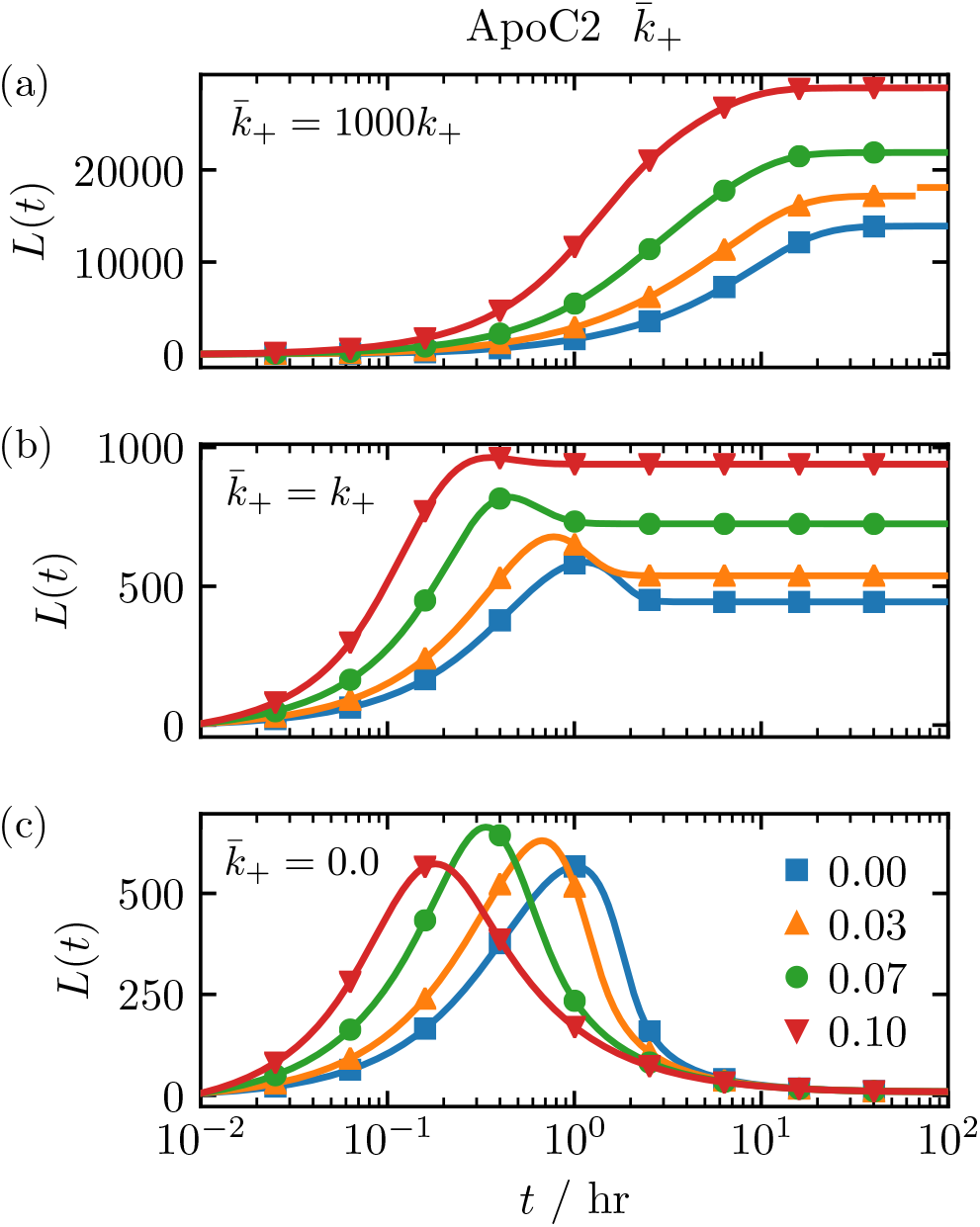
Predicted length profiles for ApoC2 aggregation. Unless otherwise noted in panels (a-c), the fit parameters used a re listed in the Supporting Information.

Finally, Fig. 6(c) shows the scenario when monomer addition dominates. At early times, clusters are sparse but grow extremely long, then monomers become starved. Fragmentation then sets in and breaks up the fibrils into a large number of much smaller clusters at later times. Increasing *ϕ* increases homogeneous cluster production at early times, and similar to actin, also reduces the maximum fibril lengths, as fewer monomers are available to join with fibrils. At later times, the steady-state lengths are longer for larger *ϕ*.

### D. Multi-step secondary nucleation

In most of the examples so far, clusters (or active polymers) were produced via homogeneous nucleation or via fragmentation. Next, we consider how crowders affect the aggregation when multi-step secondary nucleation and parallel processes contribute to cluster production. Fig. 7 shows the experimentally determined half-time dependence on *c*_0_ (blue circles) for A*β*(1-40) (Fig. 7(a)) and A*β*(1-42) (Figures 7(b) and (c)) proteins. In Fig. 7(c) the data was obtained at low ionic strengths. The red line in each plot shows the predicted relationship computed by fitting *M*(*t*)/*c*_0_ to ThT data.^49,70,71^

**FIG. 7.**
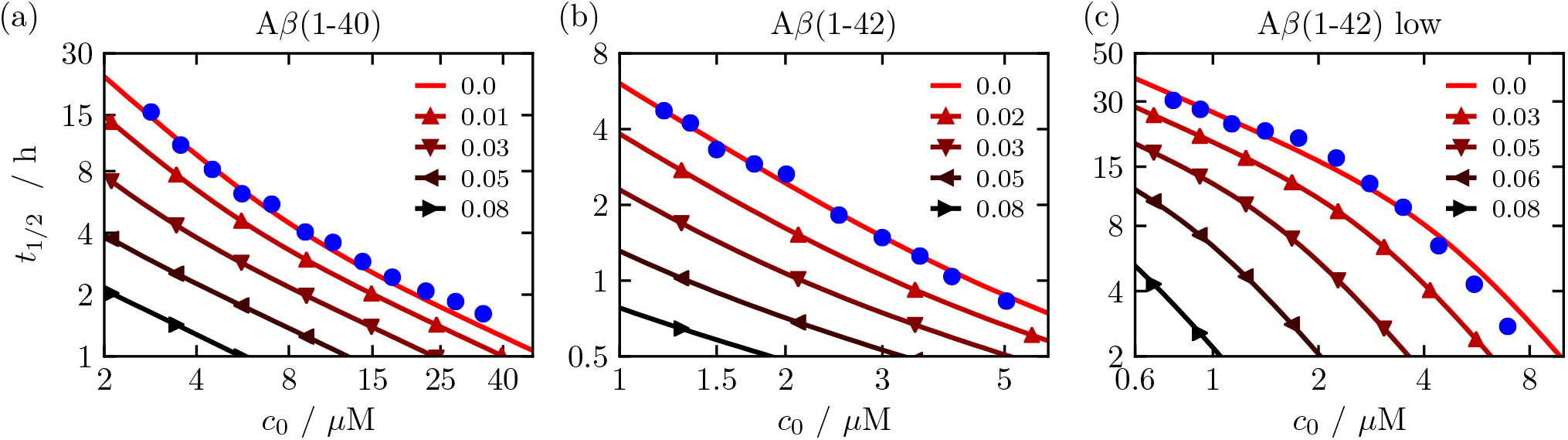
The half-time *t*_1/2_ versus initial protein monomer concentration *c*_0_ is shown for (a) A*β*(1-40), (b) A*β*(1-42), and (c) A*β*(1-42) at low ionic strengths, respectively. In each figure, blue circles show the experimental data,^49,70,71^ the red line connecting the blue circles shows the computed *M*(*t*) in a crowder-free environment, and triangles show *M*(*t*) computed for *ϕ* > 0.0.

In Fig. 7(a), aggregate production is dominated by 2-step secondary nucleation as described by Eq. (30). The positive curvature of the red curve show that the secondary nucleation is effectively 1-step at low *c*_0_ and 2-step at large *c*_0_, with the second-step saturating and becoming rate-limiting as *c*_0_ increases (note that saturation is controlled by the rate constant *K_M_*).^49^ In Fig. 7(b), a secondary nucleation mechanism also primarily drives the formation of A*β*(1-42) aggregates,^70^ but the constant slope of the red curve indicates that the low-*c*_0_ behavior (again effective 1-step secondary nucleation) is observed even at high *c*_0_. Finally, the negative curvature of the red line in Fig. 7(c) shows two parallel nucleation processes driving A*β*(1-42) aggregation at low ionic strengths.^71^ At low *c*_0_, the nucleation is mainly controlled by fragmentation while at high *c*_0_ 1-step secondary nucleation dominates as 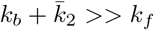 in Eq. (30).

Fig. 7 also shows the predicted scaling relationships when crowders are hypothetically introduced into the experiments. For each protein, we used the rate constants obtained from the fits to the experimental data (the red curves), and assumed a constant crowder radius of 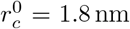. To mimic changes to the half-times seen in the experiments performed by Lee, et al.,^60^ we investigated excluded volume fractions in the same range, and used the same value for *A*_1_ and *χ* for the experiments involving agitation, while adjusting *r_sc_*, for both A*β*(1-40) and A*β*(1-42) shown in Fig. 7. Fig. 7 clearly shows for all three examples that as *ϕ* increases, the high-*ϕ* nucleation behavior dominates at progressively lower *c*_0_, where the high-*ϕ* nucleation behavior resembles the high-*c*_0_ behavior of the crowder-free case.

## VII. DISCUSSION AND CONCLUSION

In summary, we have extended a well-studied mass-action model that is based on generalized Smoluchowski kinetics to describe protein aggregation in crowded environments. This was accomplished by using SPT to take into account the changes to the activity coefficients as caused by the presence of crowders. The crowders were assumed to affect only association reactions (nucleation, monomer addition, and fibril merging). The protein monomers were also assumed to be spherical, while the protein aggregates were assumed to be sphero-cylindrical structures. Additionally, we modeled the dependency of the (effective) crowder radius, *r_c_*, on *ϕ* using a simple relationship aimed at capturing the leading order polynomial correction. This is an approximate way to take the zeroth-order effects of viscosity change due to crowders into consideration.

Overall, the model predicts that the presence of crowders can speed up, cause no change, or slow down the progress of the aggregation compared to crowder-free solutions, in excellent agreement with experimental results from nine different amyloid-forming proteins that utilized dextran as the crowder. These different dynamic effects of macromolecular crowding can be understood in terms of the change of excluded volume associated with each reaction step. The speedup is caused by the proteins taking up less excluded volume once in aggregate, thereby increasing the entropy of the crowders and lowering the barrier for proteins to join fibrils, compared to a crowder free environment. Only when the excluded volume associated with the aggregated protein is larger than that of the merging proteins do we observe a slowdown in the aggregation. This rise of excluded volume decreases the entropy of the buffer, thereby raising the freeenergy barrier for monomers or oligomers to join aggregates. Furthermore, we showed that the presence of crowders can significantly change the length distributions of the fibrils formed, and may promote either 1-step or 2-step secondary nucleation over primary nucleation.

Our results may suggest that for some proteins, a conformational change of the monomer may have occurred once incorporated into an aggregate.^69^ An increase in the number of intra-chain bonds formed, or inter-chain bonds would compact the protein compared to the monomer. The proteins shown in Fig. 3(a) may still undergo a structural rearrangement, but any changes would not cause the protein to exclude more volume on average compared with the average monomer configuration.

One other scenario that we did not consider here that could also explain the slow-down in aggregation for the proteins shown in Fig. 3(b) is the possibility for competition between monomer refolding and aggregation. This scenario was studied by Ellis and Minton, who included an unfolded/native reaction *U* ⇌ *N* in their model for describing protein aggregation kinetics.^61^ Depending on the values of the folding rate constant *k_f_*, and that for the monomer association reaction in the presence of crowders, refolding may dominate over protein aggregation, and a corresponding slow-down in the aggregation compared to a crowder-free environment could be observed. For example, the A*β*(1-40) data shown in Fig. 3 shows only the shaken environment experiencing an increase in the aggregation rate. The shaking may be preventing the monomer protein from refolding and encouraging aggregation, whereas for the unshaken system the monomers may prefer to refold rather than to aggregate.

The model does have some limitations, thus allowing for potential improvements. As noted, we did not try to account for conformational changes occurring in either monomer proteins, intermediate oligomers, or the fibrils themselves, and instead focused on polymerization mechanisms. Second, we assumed that the fibril merging and fragmentation reactions were independent of aggregate size. While this may be a physically realistic assumption for fibril fragmentation,^72^ simple size-dependent formulations of the association reaction rates based on coagulation theory predict that monomer addition should have a higher rate compared with the merging rate of two fibrils of the same size.^73^ Finally, the number of parameters needed to describe the different nucleation, growth, shrinkage, and the influence of crowders on the different rates, renders the optimization of the model parameters to the experimental data cumbersome in some cases. This problem is exasperated when solving for each *c_r_*(*t*) explicitly when size-dependent rate constants are introduced, as closed-form expressions for *P*(*t*) and *M*(*t*) cannot typically be derived.^23^

In future investigations, we plan to extend the model introduced here to describe protein conformational changes in more detail. We previously extended mass-action models based on generalized Smoluchowski kinetics to describe nucleated conformational conversion scenarios.^23^ Such a model would facilitate investigation of the competition between refolding and aggregation, nucleated conformational conversion mechanisms, as well as systems with potentially multiple aggregate species coexisting in solution that have varying levels of *β*-sheet structure. For example, the extended model could be used to investigate small oligomers, which may be initially disordered and must undergo a structural rearrangement before elongating into long fibrils.^74–76^ Additionally, the effect of adding a folding/unfolding transition for the monomer proteins to the current model will also be investigated, as well as how size-dependent fibril merging and fragmentation rates alter the bulk quantities *M*(*t*) and *L*(*t*), in future studies.

## ACKNOWLEDGMENTS

We thank Frank Ferrone for stimulating discussion on heterogeneous nucleation and associated crowding effects. J. S. S. thanks Petr Šulc for useful discussions.

