## Supporting Information for "Investigating the effects of molecular crowding on the kinetics of protein aggregation"

John S. Schreck,[1,2,a)] John Bridstrup,[3] and Jian-Min Yuan[3]
[1)] National Center for Atmospheric Research, Boulder, CO 80305
[2)] Department of Chemistry, Drexel University, Philadelphia, PA 19104
[3)] Department of Physics, Drexel University, Philadelphia, PA 19104

### SI. OBTAINING $t_{1/2}$ FROM THIOFLAVIN T FLUORESCENCE CURVES

All half-times reported in Figs. 2 and 3 were found by fitting

$$F(t) = F_0 + \frac{A}{1 + \exp\left(-k_{\text{app}}(t - t_{1/2})\right)} \tag{S1}$$

to the experimental Thioflavin T (ThT) curves as a function of time. The parameters $F_0$, $A$, $k_{\text{app}}$, and $t_{1/2}$ were adjusted to minimize the mean-average error between Eq. (S1) and the experimentally reported values. All fits of $F(t)$ to the data sets were re-scaled according to $(F(t) - \min(F(t)) / (\max(F(t)) - \min(F(t)))$ from which the half-time is obtained when this quantity equaled one half. The experimental data was similarly re-scaled using the same maximum and minimum values from the fitted $F(t)$.

### SII. RATE CONSTANTS

All non-zero fit parameters used in the figures are listed for each protein investigated. The rate constants for monomer addition and subtraction are $k_+^0$ and $k_-^0$, respectively, while we assume that merging and fragmentation mechanisms are independent of size so that $\bar{k}_+(r,s) \equiv \bar{k}_+$ and $\bar{k}_-(r,s) \equiv \bar{k}_-$.

#### A. Actin from Rosin, et al.[47]

$n_c = 3$, $m_0 = 5\,\mu\text{M}$, $k_+^0 = 455.5\,\mu\text{M}\,\text{h}^{-1}$ and $k_n^0 = 1.02 \times 10^{-5}\,\mu\text{M}^2\,\text{h}^{-1}$, $r_c^0 = 1.8\,\text{nm}$, $r_1 = r_{sc} = 2.15\,\text{nm}$, $A_1 = 2.70 \times 10^{-9}\,\text{nm}^{-\chi}$, and $\chi = 1.87$. See Fig. 2.

#### B. Human PrP from Zhou, et al.[64]

$n_c = 1$, $m_0 = 10.0\,\mu\text{M}$, $k_+ = 36.8\,\mu\text{M}\,\text{h}^{-1}$, $k_- = 0.044\,\text{h}^{-1}$, $\bar{k}_-^0 = 7.6 \times 10^{-4}\,\text{h}^{-1}$, $r_c^0 = 1.8\,\text{nm}$, $r_1 = r_{sc} = 3.5\,\text{nm}$, $A_1 = 2.70 \times 10^{-9}\,\text{nm}^{-\chi}$, $\chi = 1.87$. See Fig. S1.

#### C. ApoC2 from Binger, et al.[?]

$n_c = 5$, $m_0 = 45\,\mu\text{M}$, $k_+^0 = 24.49\,\mu\text{M}\,\text{h}^{-1}$, $k_-^0 = 1.48\,\text{h}^{-1}$, $\bar{k}_+^0 = 12.96 \times 10^3\,\mu\text{M}\,\text{h}^{-1}$, $\bar{k}_-^0 = 5.70 \times 10^{-3}\,\text{h}^{-1}$, $k_n^0 = 1.54 \times 10^{-11}\,\mu\text{M}^4\,\text{h}^{-1}$,$r_c^0 = 1.24\,\text{nm}$, $r_1 = r_{sc} = 2.10\,\text{nm}$, $A_1 = 6.10 \times 10^{-10}\,\text{nm}^{-\chi}$, and $\chi = 1.95$. See Fig. S2.

#### D. $A\beta$(1-40) from Lee, et al.[60]

$n_c = n_2 = 2$, $m_0 = 30.0\,\mu\text{M}$, $k_+^0 = 72.05\,\mu\text{M}\,\text{h}^{-1}$, $k_-^0 = 0.0033\,\text{h}^{-1}$, $\bar{k}_+^0 = 3.07 \times 10^{-4}\,\mu\text{M}\,\text{h}^{-1}$, $\bar{k}_-^0 = 3.54 \times 10^{-3}\,\text{h}^{-1}$, $k_n^0 = 1.13 \times 10^{-19}\,\mu\text{M}\,\text{h}^{-1}$, $\bar{k}_2^0 = 7.47 \times 10^{-10}\,\text{h}^{-1}$, $k_f = 1.31\,\text{h}^{-1}$, $k_b = 0.26\,\text{h}^{-1}$, $r_c^0 = 1.8\,\text{nm}$, $r_1 = 2.26\,\text{nm}$, $r_{sc} = 12.3\,\text{nm}$, $A_1 = 2.35 \times 10^{-8}\,\text{nm}^{-\chi}$, and $\chi = 0.67$. See Fig. S3.

#### E. $\beta$-LAC from Ma, et al.[66]

$n_c = 4$, $m_0 = 82\,\mu\text{M}$, $k_+^0 = 1.04 \times 10^3\,\mu\text{M}\,\text{h}^{-1}$, $k_-^0 = 7.68 \times 10^{-7}\,\text{h}^{-1}$, $\bar{k}_+^0 = 2.05 \times 10^{-3}\,\text{h}^{-1}$, $\bar{k}_-^0 = 9.76 \times 10^{-9}\,\text{h}^{-1}$, $k_n^0 = 4.10 \times 10^{-22}\,\mu\text{M}^3\,\text{h}^{-1}$, $r_c^0 = 1.8\,\text{nm}$, $r_1 = r_{sc} = 2.01\,\text{nm}$, $A_1 = 2.77 \times 10^{-9}\,\text{nm}^{-\chi}$, and $\chi = 1.94$. See Fig. S4.

#### F. $\beta$2m from Luo, et al.[65]

$n_c = 1$, $m_0 = 1.4\,\mu\text{M}$, $k_+^0 = 1.08 \times 10^2\,\mu\text{M}\,\text{h}^{-1}$, $k_-^0 = 1.2 \times 10^{-6}\,\text{h}^{-1}$, $\bar{k}_-^0 = 4.8 \times 10^{-3}\,\text{h}^{-1}$, $r_c^0 = 1.8\,\text{nm}$, $r_1 = r_{sc} = 2.74\,\text{nm}$, $A_1 = 2.92 \times 10^{-9}\,\text{nm}^{-\chi}$, and $\chi = 1.82$. See Fig. S5.

#### G. $A\beta$(1-40) from Meisl, et al.[49]

$n_c = n_2 = 2$, $m_0 = 5.0\,\mu\text{M}$, $k_+^0 = 3.60\,\mu\text{M}\,\text{h}^{-1}$, $k_n^0 = 1.19 \times 10^{-5}\,\mu\text{M}\,\text{h}^{-1}$, $\bar{k}_2^0 = 6.30 \times 10^{-2}\,\text{h}^{-1}$, $k_f = 0.63\,\text{h}^{-1}$, $k_b = 30.38\,\text{h}^{-1}$, $r_c^0 = 1.8\,\text{nm}$, $r_1 = 2.26\,\text{nm}$, $r_{sc} = 2.4\ r_1$, $A_1 = 2.35 \times 10^{-8}\,\text{nm}^{-\chi}$, and $\chi = 0.67$. See Fig. 7(a).

#### H. $A\beta$(1-42) from Cohen, et al.[70]

$n_c = n_2 = 2$, $m_0 = 5.0\,\mu\text{M}$, $k_+^0 = 85.0\,\mu\text{M}\,\text{h}^{-1}$, $k_n^0 = 1.72 \times 10^{-6}\,\mu\text{M}\,\text{h}^{-1}$, $\bar{k}_2^0 = 2.76 \times 10^{-1}\,\text{h}^{-1}$, $k_f = 8.99 \times 10^{-2}\,\text{h}^{-1}$, $k_b = 1.24\,\text{h}^{-1}$, $r_c^0 = 1.8\,\text{nm}$, $r_1 = 2.26\,\text{nm}$, $r_{sc} = 2.0\ r_1$, $A_1 = 2.35 \times 10^{-8}\,\text{nm}^{-\chi}$, and $\chi = 0.67$. See Fig. 7(b).

[a)] Electronic mail:

#### I. $A\beta$(1-42)-low from Meisl, et al.[71]

$n_c = n_2 = 3$, $m_0 = 5.0\,\mu\text{M}$, $k_+^0 = 8.34\,\mu\text{M}\,\text{h}^{-1}$, $\bar{k}_-^0 = 1.43 \times 10^{-2}\,\text{h}^{-1}$, $k_n^0 = 9.91 \times 10^{-8}\,\mu\text{M}\,\text{h}^{-1}$, $\bar{k}_2^0 = 1.74 \times 10^3\,\text{h}^{-1}$, $k_f = 1.72 \times 10^2\,\text{h}^{-1}$, $k_b = 1.78 \times 10^9\,\text{h}^{-1}$, $r_c^0 = 1.8\,\text{nm}$, $r_1 = 2.26\,\text{nm}$, $r_{sc} = 2.5\,r_1$, $A_1 = 2.35 \times 10^{-8}\,\text{nm}^{-\chi}$, and $\chi = 0.67$. See Fig. 7(c).

#### J. IAPP from Seeliger, et al.[67]

$n_c = 1$, $m_0 = 10.0\,\mu\text{M}$, $k_+^0 = 20.54\,\mu\text{M}\,\text{h}^{-1}$, $k_-^0 = 5.11 \times 10^{-2}\,\text{h}^{-1}$, $\bar{k}_-^0 = 2.28 \times 10^{-3}\,\text{h}^{-1}$, $k_n^0 = 4.86 \times 10^{-4}\,\mu\text{M}\,\text{h}^{-1}$, $r_c^0 = 1.8\,\text{nm}$, $r_1 = r_{sc} = 2.74\,\text{nm}$, $A_1 = 2.92 \times 10^{-9}\,\text{nm}^{-\chi}$, and $\chi = 1.82$. See Fig. S6.

#### K. Rabbit PrP from Zhou, et al.[64]

$n_c = 1$, $m_0 = 10.0\,\mu\text{M}$, $k_+^0 = 36.83\,\mu\text{M}\,\text{h}^{-1}$, $k_-^0 = 4.41 \times 10^{-2}\,\text{h}^{-1}$, $\bar{k}_+^0 = 6.56 \times 10^{-4}\,\mu\text{M}\,\text{h}^{-1}$, $\bar{k}_-^0 = 2.32 \times 10^{-3}\,\text{h}^{-1}$, $k_n^0 = 1.86 \times 10^{-6}\,\mu\text{M}^4\,\text{h}^{-1}$, $r_c^0 = 1.8\,\text{nm}$, $r_1 = 3.34\,\text{nm}$, $r_{sc} = 2.21\,\text{nm}$ $A_1 = 4.65 \times 10^{-11}\,\text{nm}^{-\chi}$, and $\chi = 2.04$. See Fig. S7.

#### L. BCA from Mittal, et al.[68]

$n_c = 1$, $m_0 = 7.0\,\mu\text{M}$, $k_+^0 = 1161.56\,\mu\text{M}\,\text{h}^{-1}$, $k_-^0 = 48.31\,\text{h}^{-1}$, $\bar{k}_-^0 = 9.45 \times 10^{-2}\,\text{h}^{-1}$, $k_n^0 = 4.14 \times 10^{-3}\,\mu\text{M}^4\,\text{h}^{-1}$, $r_c^0 = 1.8\,\text{nm}$, $r_1 = 3.16\,\text{nm}$, $r_{sc} = 2.27\,\text{nm}$ $A_1 = 4.72 \times 10^{-10}\,\text{nm}^{-\chi}$, and $\chi = 1.84$. See Fig. S8.

#### M. $A\beta$(1-40) from Lee, et al.[60]

$n_c = n_2 = 2$, $m_0 = 30.0\,\mu\text{M}$, $k_+^0 = 7.50\,\mu\text{M}\,\text{h}^{-1}$, $k_-^0 = 6.59 \times 10^{-11}\,\text{h}^{-1}$, $\bar{k}_-^0 = 1.58 \times 10^{-9}\,\text{h}^{-1}$, $k_n^0 = 8.39 \times 10^{-9}\,\mu\text{M}\,\text{h}^{-1}$, $\bar{k}_2^0 = 1.28 \times 10^{-5}\,\text{h}^{-1}$, $k_f = 5.09 \times 10^8\,\text{h}^{-1}$, $k_b = 2.6 \times 10^{10}\,\text{h}^{-1}$, $r_c^0 = 1.8\,\text{nm}$, $r_1 = 2.41\,\text{nm}$, $r_{sc} = 1.56\,\text{nm}$, $A_1 = 1.26 \times 10^{-9}\,\text{nm}^{-\chi}$, and $\chi = 0.50$. See Fig. S9.

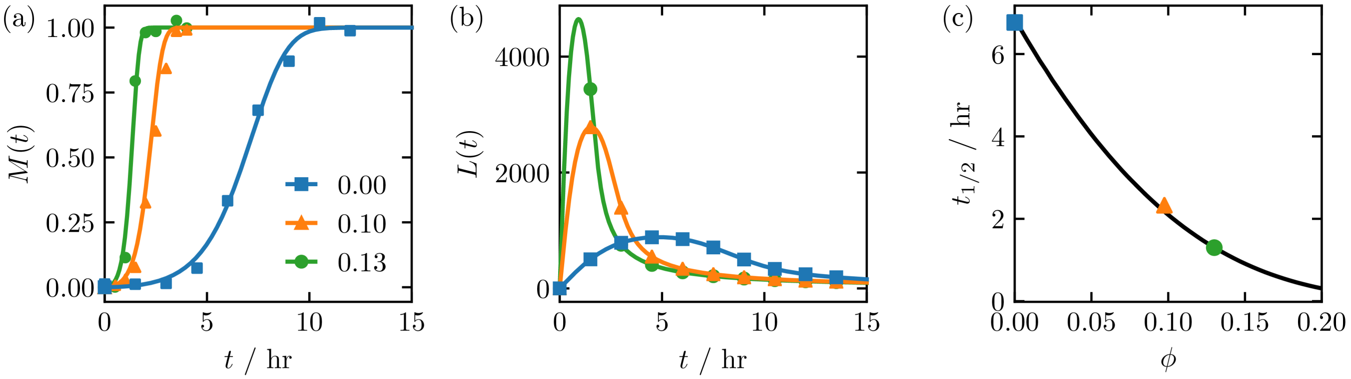


FIG. S1. Human PrP. In (a) the mass concentration of fibrils $M(t)$ is shown and was fit to ThT data from Zhou, et al.[64] In (b) the predicted $L(t)$ is shown for different crowder concentrations $\phi$. In (c) the half-time ($t_{1/2}$) obtained from the curves in (a) is shown versus $\phi$.The different symbol shapes indicate different values for $\phi$. In (a) they also refer to experimental data.

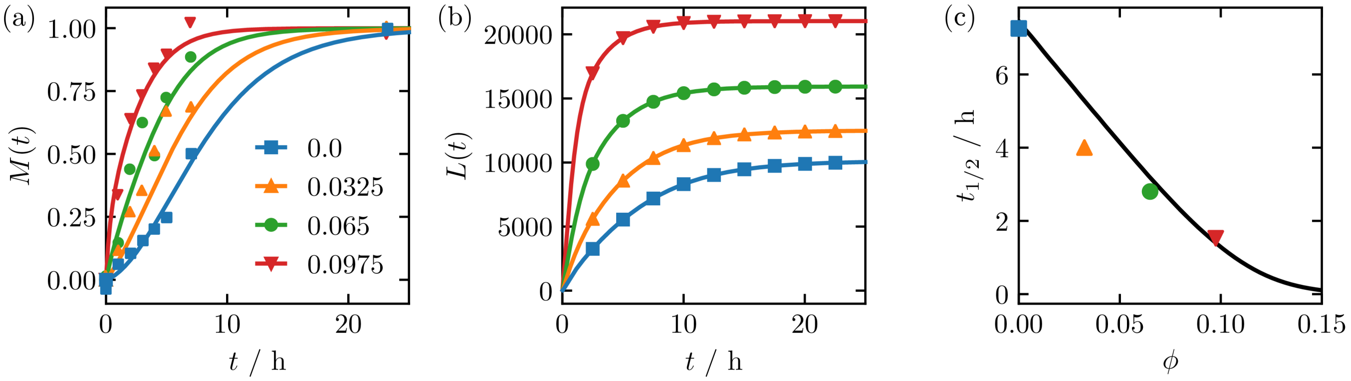


FIG. S2. ApoC2. In (a) the mass concentration of fibrils $M(t)$ is shown and was fit to ThT data from Binger, et al.[?] In (b) the predicted $L(t)$ is shown for different crowder concentrations $\phi$. In (c) the half-time ($t_{1/2}$) obtained from the curves in (a) is shown versus $\phi$.The different symbol shapes indicate different values for $\phi$. In (a) they also refer to experimental data.

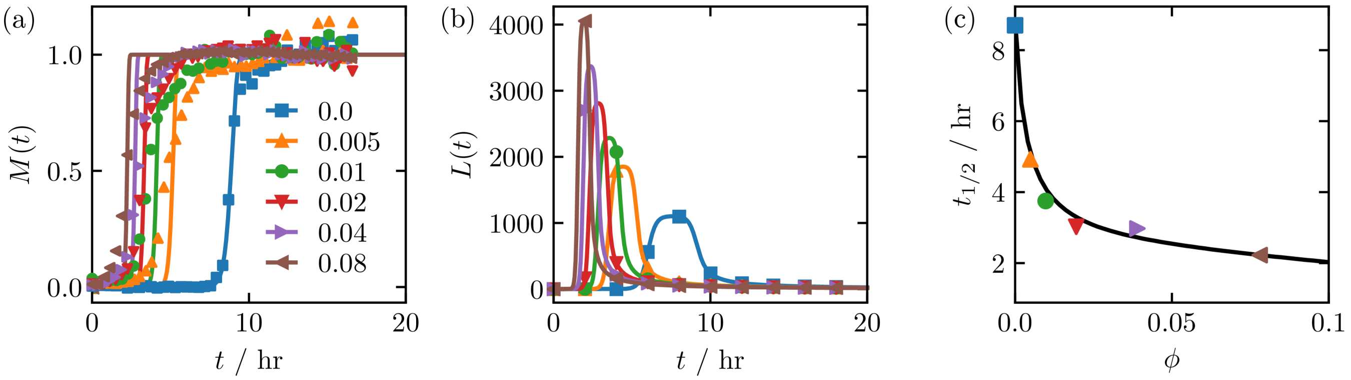


FIG. S3. $A\beta$(1-40) in agitated (shaken) conditions. In (a) the mass concentration of fibrils $M(t)$ is shown and was fit to ThT data from Lee, et al.[60] In (b) the predicted $L(t)$ is shown for different crowder concentrations $\phi$. In (c) the half-time ($t_{1/2}$) obtained from the curves in (a) is shown versus $\phi$.The different symbol shapes indicate different values for $\phi$. In (a) they also refer to experimental data.

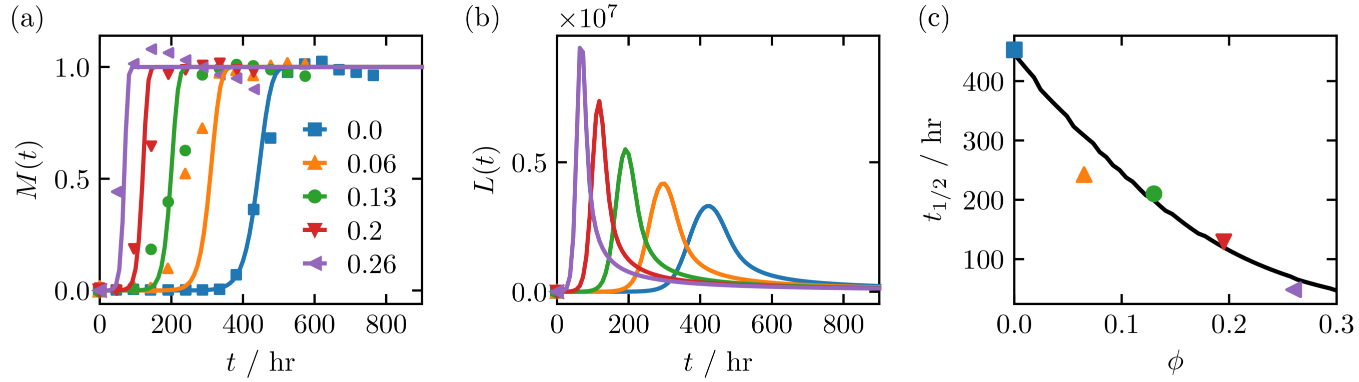


FIG. S4. $\beta$-LAC. In (a) the mass concentration of fibrils $M(t)$ is shown and was fit to ThT data from Ma, et al.[66] In (b) the predicted $L(t)$ is shown for different crowder concentrations $\phi$. In (c) the half-time ($t_{1/2}$) obtained from the curves in (a) is shown versus $\phi$.The different symbol shapes indicate different values for $\phi$. In (a) they also refer to experimental data.

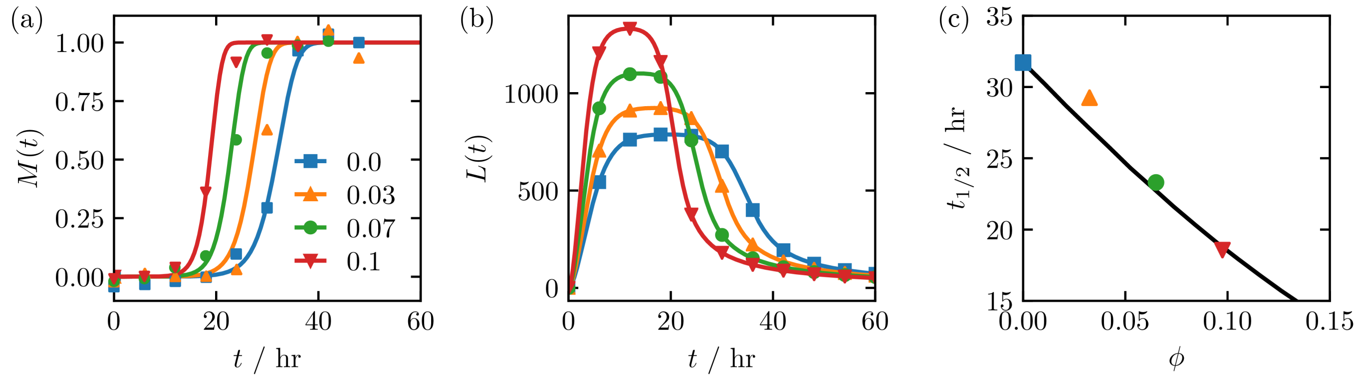


FIG. S5. $\beta$2m. In (a) the mass concentration of fibrils $M(t)$ is shown and was fit to ThT data from Luo, et al.[65] In (b) the predicted $L(t)$ is shown for different crowder concentrations $\phi$. In (c) the half-time ($t_{1/2}$) obtained from the curves in (a) is shown versus $\phi$.The different symbol shapes indicate different values for $\phi$. In (a) they also refer to experimental data.

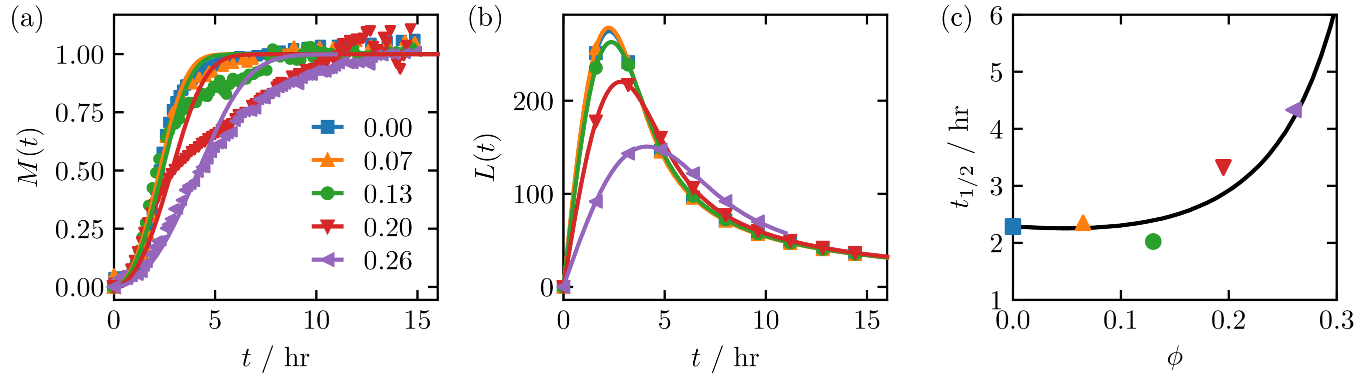


FIG. S6. IAPP. In (a) the mass concentration of fibrils $M(t)$ is shown and was fit to ThT data from Seeliger, et al.[67] In (b) the predicted $L(t)$ is shown for different crowder concentrations $\phi$. In (c) the half-time ($t_{1/2}$) obtained from the curves in (a) is shown versus $\phi$.The different symbol shapes indicate different values for $\phi$. In (a) they also refer to experimental data.

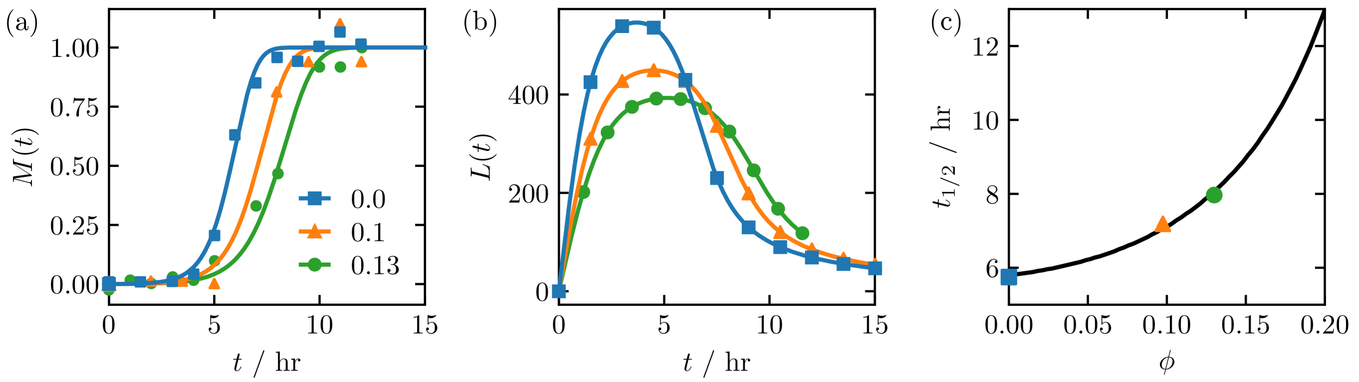


FIG. S7. Rabbit PrP. In (a) the mass concentration of fibrils $M(t)$ is shown and was fit to ThT data from Zhou, et al.[64] In (b) the predicted $L(t)$ is shown for different crowder concentrations $\phi$. In (c) the half-time ($t_{1/2}$) obtained from the curves in (a) is shown versus $\phi$.The different symbol shapes indicate different values for $\phi$. In (a) they also refer to experimental data.

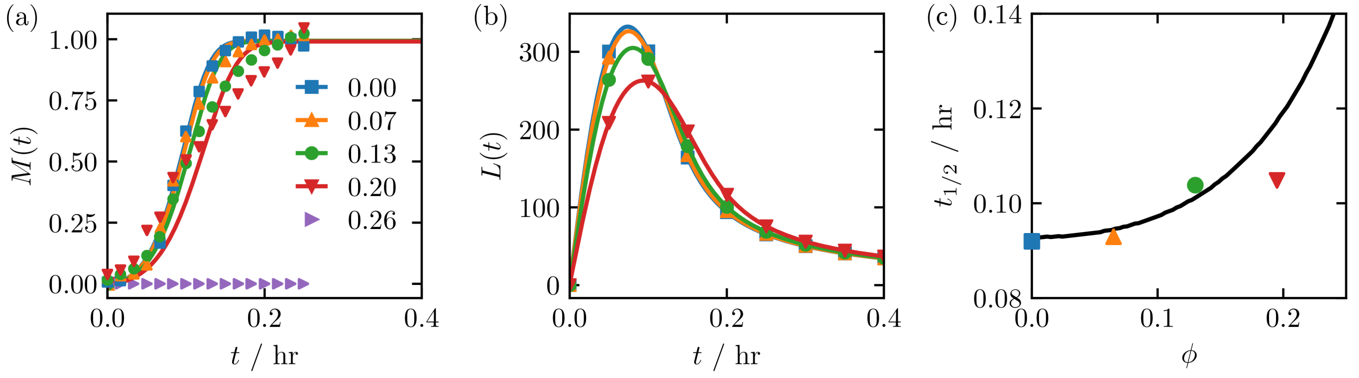


FIG. S8. BCA. In (a) the mass concentration of fibrils $M(t)$ is shown and was fit to ThT data from Mittal, et al.[68] We could not obtain the half-time for the data at $\phi = 0.26$ (purple right-facing triangles) using Eq. (S1). In (b) the predicted $L(t)$ is shown for different crowder concentrations $\phi$. In (c) the half-time ($t_{1/2}$) obtained from the curves in (a) is shown versus $\phi$.The different symbol shapes indicate different values for $\phi$. In (a) they also refer to experimental data.

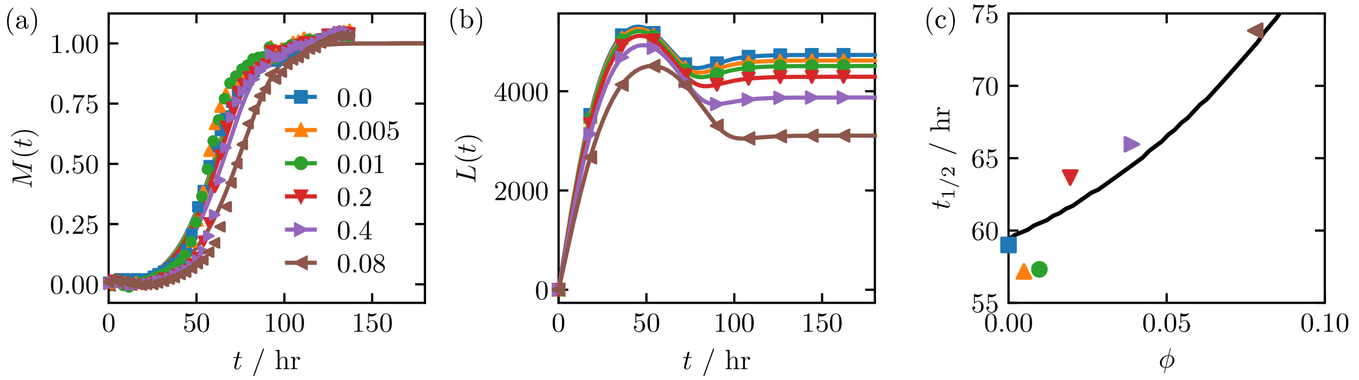


FIG. S9. $A\beta$(1-40). In (a) the mass concentration of fibrils $M(t)$ is shown and was fit to ThT data from Lee, et al.[60] In (b) the predicted $L(t)$ is shown for different crowder concentrations $\phi$. In (c) the half-time ($t_{1/2}$) obtained from the curves in (a) is shown versus $\phi$.The different symbol shapes indicate different values for $\phi$. In (a) they also refer to experimental data.
